# Gut microbiome composition predicts summer core range size in a generalist and specialist ungulate

**DOI:** 10.1101/2020.08.04.236638

**Authors:** J.F. Wolf, K.D. Kriss, K.M. MacAulay, K. Munro, B.R. Patterson, A.B.A. Shafer

## Abstract

The gut microbiome of animals varies by age, diet, and habitat, and directly influences individual health. Similarly, variation in an individual’s home range can lead to differences in feeding strategies and fitness. Ungulates (hooved mammals) exhibit species-specific microbiomes and habitat use patterns: here, we combined gut microbiome and movement data to assess relationships between space use and the gut microbiome in a specialist and a generalist ungulate. We captured and GPS radiocollared 24 mountain goats (*Oreamnos americanus*) and 34 white-tailed deer (*Odocoileus virginianus*). We collected fecal samples and conducted high-throughput sequencing of the 16S rRNA gene. Using GPS data, we estimated core (50%) and home range (95%) sizes and calculated proportional use for several important habitat types. We generated metrics related to gut diversity and key bacterial ratios. We hypothesize that larger *Firmicutes* to *Bacteroides* ratios confer body size or fat advantages that allow for larger home ranges, and that relationships between gut diversity and disproportionate habitat use is stronger in mountain goats due to their restricted niche relative to white-tailed deer. *Firmicutes* to *Bacteroides* ratios were positively correlated with core range area in both species. Mountain goats exhibited a negative relationship between gut diversity and use of two key habitat types (treed areas and escape terrain), whereas no relationships were detected in white-tailed deer. This is the first study to relate core range size to the gut microbiome in wild ungulates and is an important proof of concept that advances the information that can be gleaned from non-invasive sampling.

## Introduction

The gastrointestinal tract of animals contains trillions of microbes that influence individual health. Gut bacteria, hereafter termed the gut microbiome, can modify immune responses [1], improve and modulate metabolism [2], and affect behaviour [3,4]. While largely stable over time [5,6], disturbance of the gut microbiome can lead to disease [7] and impacts metabolic versatility, meaning the ability to survive equally well when presented with a wide range of dietary compositions and habitat [8,9]. Gut microbiome diversity has been shown to impact behaviour; for example, gut microbiome manipulation in mice resulted in improved memory as measured using a passive-avoidance test [10]. In another example, the presence of specific gut bacteria species suppressed protein appetite, indicating the ability of the gut microbiome to drive dietary decisions [11]. Differences in gut microbiome composition are correlated with landscape alteration [12,13], while levels of daily activity and foraging also appear to be influenced by the gut microbiome [14,15]. Notably, distinct diet types, such as herbivory and carnivory, are associated with unique microbiome profiles [16]. The mechanistic links are not totally understood, but are thought to follow the microbiota-gut-brain axis where bacteria have the ability to generate neurotransmitters that influence cognition [17].

Mammalian herbivores are typified by specific gut microbial taxa as they rely on these bacteria to extract energy and nutrients from food, synthesize vitamins, and detoxify plant defense compounds [18]. For example, *Firmicutes* and *Bacteroidetes* are involved in energy resorption and carbohydrate metabolism; *Firmicutes* can act as a more effective energy source, leading to more efficient calorie absorption and weight gain, while *Bacteroidetes* are involved with energy production and conversion, and amino acid transport and metabolism [19]. Ungulates, and ruminants in particular, have specialized anatomical and physiological adaptations to accommodate the cellulolytic fermentation of low-nutrition, high-fiber plant materials [20]. A specialized gut microbiome allows ruminants to digest typically indigestible plant biomass [21] and as a result, exploit novel environments. Mountain goats (*Oreamnos americanus*) are alpine ruminants that are endemic to the mountainous regions of northwestern North America [22]. Mountain goats use lower elevation, forested, and warmer aspect habitat during the winter and higher elevation, mountainous terrain in summer [23–26]. They are considered intermediate browsers and eat a variety of forage, with diets generally dominated by grasses [27,28]. In contrast, white-tailed deer (*Odocoileus virginianus*) exploit a variety of habitat and food resources and cover a large geographic range that stretches across most of North America and includes part of Central and South America [29]. White-tailed deer use woody cover habitats year-round, but can also thrive in urban and rural settings [30,31]; they maintain distinct seasonal ranges in the northern parts of their range and are considered browsing ruminants and both habitat and dietary generalists [32].

Our study integrated high-throughput sequencing and GPS telemetry to evaluate the relationship between gut microbiome, home range area, and disproportionate habitat use of two ungulates living in contrasting environments. This is among the first studies to link space use to the gut microbiome in wild systems and provides a proof of concept that advances the information that can be generated from non-invasive sampling. We quantified the relationship between key microbiome diversity metrics on home range size and relative use of different habitat classes inferred from GPS tracking of individuals. From an evolutionary perspective, this link between variation in phenotype or behaviour and the gut microbiome assumes selection operates on both the phenotypes of the constituents (microbiome) and host, with the collective genetic material known as the hologenome [33]. We hypothesized that an increase in gut diversity would be linked with an increase in area used, as greater gut diversity would reflect, and possibly drive larger use of space and a more resource-diverse home range, similar to the findings of [34]. As high *Firmicutes* to *Bacteroides* ratios correspond to larger body size and fat stores [19,35], we hypothesized that larger *Firmicutes* to *Bacteroides* ratios would be correlated with larger home ranges, as individuals building up fat stores for winter would generally use more space to forage. This relationship may be impacted by resource distribution, as specialists prefer homogenously distributed resources, while generalists prefer heterogeneously distributed resources, which can impact space use [36]. Consequently, we hypothesized that relationships between proportional habitat use and the gut microbiome would be stronger in specialists as they have a more restricted niche with deviations from this having larger consequences, whereas generalists can make use of a variety of habitat areas.

## Methods

### Animal captures, sample collection and DNA extraction

We captured and radio-collared male and female mountain goats on three adjacent mountain complexes using aerial net-gun capture northeast of Smithers, British Columbia, Canada (Fig. 1). We captured and radio-collared 34 female white-tailed deer using baited Clover traps southwest of Ottawa, Ontario, Canada (Fig. 1). For more information on animal captures, see [37,38]. VERTEX Plus and VERTEX Lite Global Positioning System (GPS) collars (VECTRONIC Aerospace, Germany) were used for mountain goats, while store-on-board (G2110D, Advanced Telemetry Solutions, Isanti, MN) or GSM-upload (Wildcell SG, Lotek Wireless Wildlife Monitoring, Newmarket, ON) GPS collars were used for white-tailed deer. Collars recorded locations every four hours for mountain goats and every five hours for white-tailed deer. During captures we took fresh fecal pellets from each individual and stored them at −20°C; all captures took place during winter. Lab surfaces were sterilized with 90% EtOH and 10% bleach solution and a small portion of a single fecal sample (~1/4 including exterior and interior portions) was digested overnight at 56°C in 20 ul proteinase K and 180 ul Buffer ATL from the Qiagen DNeasy Blood & Tissue Kit (Qiagen, Valencia, California, USA). DNA was extracted from the digest with the QIAamp PowerFecal DNA Kit (Qiagen, Valencia, California, USA).

**Fig. 1.**
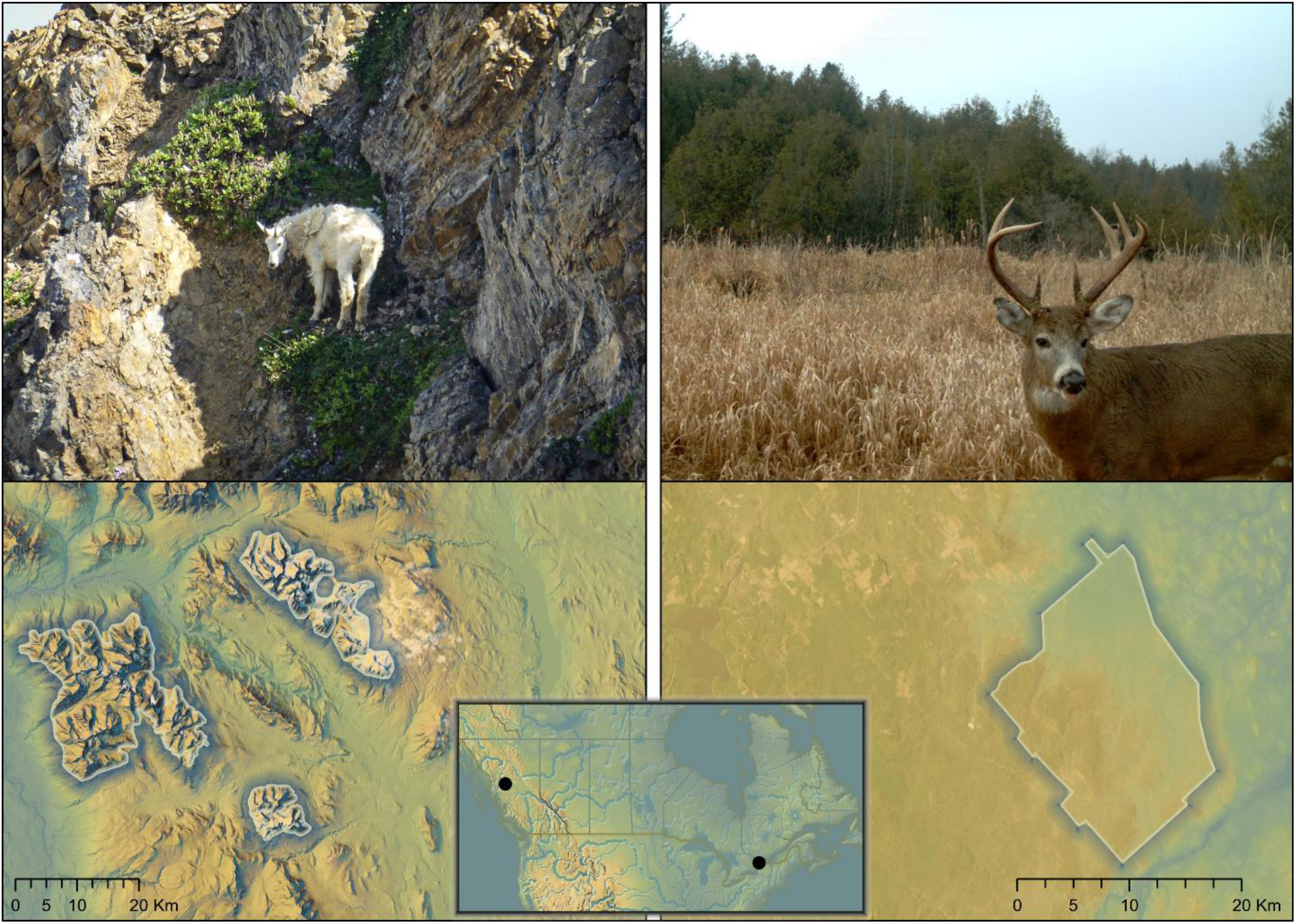
An example of habitat of both mountain goat (*Oreamnos americanus*) and white-tailed deer (*Odocoileus virginianus*) along with inset sample maps. Colors are reflective of relative elevation and topography. Average elevation of mountain goat habitat was 1614.7m, relative to 113.3m for white-tailed deer.

### High-throughput sequencing and bioinformatics

The validated Illumina 16S rRNA Metagenomic Sequencing Library Preparation (#15044223 rev. B) protocol was followed for library preparation using slight modifications [39]. The V3 and V4 regions of the 16S ribosomal ribonucleic acid (16S rRNA) hypervariable region were targeted with four variants of 341F and 805R primers designed by [40]. A unique combination of Nextera XT indexes, index 1 (i7), and index 2 (i5) adapters were assigned to each sample for multiplexing and pooling. Four replicates of each sample of fecal DNA were amplified in 25 μl PCR using the 341F and 805R primers. The replicated amplicons for each sample were combined into a single reaction of 100 μl and purified using a QIAquick PCR Purification Kit (Qiagen, 28104) and quantified on the Qubit Fluorometer. Sample indexes were annealed to the amplicons using an 8-cycle PCR reaction to produce fragments approximately 630 bp in length that included ligated adaptors; the target amplicon is approximately 430 bp in length (Illumina 16S rRNA Metagenomic Sequencing Library Preparation; #15044223 rev. B). Samples were purified with the QIAquick PCR Purification Kit and the final purified library was validated on a TapeStation (Agilent, G2991AA) and sequenced in 300 bp pair-end reads on an Illumina MiSeq sequencer at the Genomic Facility at the University of Guelph (Guelph, Ontario).

The quality of the raw sequences was assessed using FastQC v 0.11.9 (https://github.com/s-andrews/FastQC) and we determined the low-quality cut-off for forward and reverse reads. Forward and reverse reads were imported into QIIME2 v 2019.4 [41] for quality control, sequence classification, and diversity analysis. Merged, forward, and reverse reads were analyzed independently using the quality control function within QIIME2 and DADA2 to perform denoising and to detect and remove chimeras. QIIME2 follows the curated DADA2 R library workflow (https://benjjneb.github.io/dada2/) that requires zero mismatches in overlapping reads for successful merging, since reads are denoised and errors are removed before merging occurs. The taxonomy, to the species level, of all sample reads were assigned using the Silva 132 reference taxonomy database (https://docs.qiime2.org/2019.4/data-resources/). We calculated the relative proportion of *Firmicutes* to *Bacteroidetes* for each of the grouped data. Estimates of diversity included Shannon’s Index, observed richness, measured with Amplicon Sequence Variants (ASVs), and Pielou’s evenness, a measure of diversity that is the ratio of observed diversity to the maximum possible in a sample having the same number of species [42]; these were screened for correlation to one-another and read-depth. Pielou’s evenness, observed richness (ASVs), and the ratio of *Firmicutes* to *Bacteroidetes* were retained for subsequent analyses. We rarefied sample reads to the sample with the least number of reads. A principle coordinates analysis (PCoA) was also conducted using Bray Curtis dissimilarity matrices between species to determine if gut microbiome communities were different between hosts.

### GPS filtering, home range, and proportional habitat use analysis

We used different filtering approaches and seasonal delineations for each species due to the differences in landscapes occupied by mountain goats and white-tailed deer. For the mountain goat data, any N.A. or mortality signals were filtered out as were any GPS points outside of 600m - 2500m in elevation, as this reflects the maximum and minimum for the study area. Dilution of precision (DOP) values over 10 were plotted against elevation and landscape type, to ensure there were no patterns in distribution, and that filtering would not bias downstream analyses. Movement rates between successive GPS points were also calculated, and any movement rates beyond 15 km/hr were removed from analyses as these were deemed spurious. Seasons were defined as follows: Summer - May 1^st^ to October 31^st^ and Winter - December 1^st^ to April 30^th^ [23,43–45]. November was excluded from seasonal data as [26] noted a large increase in male mountain goat home ranges due to the rut. In white-tailed deer filtering occurred as above, but with no elevation restrictions as the topography of Marlborough Forest is effectively flat. White-tailed deer in Marlborough Forest have distinct summer and winter ranges; as considerable variation in migration dates were observed, movements to and from Marlborough forest were relative to each individual deer’s movement and as such, there was no hard date range. A migration movement was defined as when a deer moved between non-overlapping seasonal ranges and then occupied one seasonal range until the following migration movement [38]. No seasonal GPS data were excluded for white-tailed deer, as changes in movement patterns and home range size during the rut are minimal in females [46].

We used Brownian Bridge Movement Models (BBMMs) to generate individual home ranges using the BBMM package in R [47]. A BBMM is a continuous-time stochastic movement model that uses probabilistic and maximum likelihood approaches where observed locations are measured with error to model home ranges [48]. A minimum of 275 GPS points was required to generate a BBMM and individual home ranges were calculated with a maximum lag time between successive locations of two times the expected fix rate. A location error of 20 m was used as per [49], with a cell size of 25 m^2^. We generated 50% and 95% isopleths representing the core and home ranges respectively. Isopleths were generated separately for each individual during both summer and winter, as ungulates exhibit sex-specific habitat use patterns, that vary both seasonally and by age-class [22,26,44,50,51]. We focused on summer isopleths for both species in downstream analyses, as deer were baited in winter, which has been shown to bias movement and shift core ranges [52]. We report only the 50% isopleths, hereafter termed core ranges, to maximize seasonal differences, as they were highly correlated to the 95% isopleths (t_46_=9.3, r=0.81, p<0.0001) and results were similar between summer 50% and 95% isopleths (Fig. S1). We generated BBMM isopleths using the R package rgdal [53] and all analyses were conducted in R v.3.6.1. We imported shapefiles into ArcGIS Pro 2.5.0 to calculate home and core range areas in km^2^ for further analyses.

Proportional use of habitats was assessed by calculating the number of GPS points in a given habitat type within an isopleth, relative to the total number of GPS points located in that isopleth. Similar to Johnson’s third-order habitat selection, proportional habitat use in this study refers to how specific habitat types were used within a core range [54]. Proportion values in the 50% isopleths were highly correlated to the 95%isopleths (t117=106, r=0.99, p <2.2e-16, Table S1, S2). We selected ecologically relevant features that showed previous evidence for use in both species. Features used in mountain goat models were treed habitat (landscape classified as sub-boreal spruce, interior cedar, or Engelmann spruce), Heat Load Index (HLI; a combination of latitude, aspect, and slope), and escape terrain (landscape where slope is ≥ 40°; [55]). For HLI [56], the average value of HLI for all GPS points within an isopleth for a given individual was calculated. These features have shown evidence for disproportionate usage/selection in previous research on mountain goats [55]. We used the Biogeoclimactic Ecosystem Classification (BEC) dataset from GeoBC to determine terrestrial landcover type for mountain goats and the Digital Elevation Model from GeoBC to calculate escape terrain. In the white-tailed deer models, we used forested habitat (tree cover >60%), treed swamp (tree cover >25%), and thicket swamp (tree cover <25%, open and shrub communities), as each of the three habitat features exhibited >20% core landcover composition in [38], and thus, were available for the majority of individuals to use [57]. The Southern Ontario Land Resource Information System (SOLRIS) data set version 2.0 [58] was used to determine land cover types for white-tailed deer.

### Generalized linear models

We analyzed the associations between core range size, gut microbiome metrics, and age class for both species individually using Generalized Linear Models (GLMs) with the Gaussian family distribution and identity link function. The core range GLMs consisted of core range size as a response variable, a single microbiome metric (*Firmicutes* to *Bacteroidetes* ratio, number of ASVs, or Pielou’s evenness) and age class (adult or subadult) as fixed explanatory effects. The proportional habitat use GLMs consisted of proportion of habitat used as a response variable, a single microbiome metric (*Firmicutes* to *Bacteroidetes* ratio, ASV, or Pielou’s evenness) and age class, as fixed explanatory effects. One exception to this was the HLI GLM, as the response variable was the mean HLI value for GPS points in the isopleth, while the explanatory variables were the same as described above. Individuals 0 - 2 years of age were considered subadults for white-tailed deer, while individuals 0 - 3 years of age were considered subadults for mountain goats [22,59]. Sex was included only in mountain goat models as we only had data from female deer. Effect size and confidence intervals are reported for each model. We conducted five-fold linear model cross validation using the Caret package in R [60] to test for overfitting of our models and quantify the model’s predictive ability. We reported the Scatter Index (SI) and Root Mean Square Error (RMSE): low values in RMSE and SI are indicative of a good model fit and low residual variance.

## Results

### Bioinformatic filtering and taxonomic analysis

Twenty-three mountain goat and twenty-five white-tailed deer fecal sample sequences passed QC and a total of ~8.17 million paired-end reads (n_MG_=5,488,856, n_WTD_=2,679,668) were generated (SRA accession number PRJNA638162). FastQC analysis indicated that both forward and reverse reads lost quality > 259 bp in length (Phred score <25), so all reads were trimmed to a length of 259 bp. Following DADA2 strict quality filtering, ~1.16 million paired-end reads (n_MG_=709,457, n_WTD_=457,541) were retained for taxonomic and diversity analyses. Losing this many reads to quality filtering is typical (see [39]), as permitted error rates are extremely low in DADA2, resulting in high certainty among retained reads [61]. White-tailed deer had higher averages of both Pielou’s evenness (mean 0.95, min 0.92, max 0.96, SD 0.012) and *Firmicutes* to *Bacteroidetes* ratios (mean 8.3, min 1.49, max 21.5, SD 6.22) than mountain goats (Pielou’s evenness mean 0.92, min 0.84, max 0.95, SD 0.028; *Firmicutes* to *Bacteroidetes* ratio mean 6.90, min 3.51, max 12.10, SD 2.43). Conversely, mountain goats had higher averages of ASVs (mean 31,393.6, min 6638, max 59,481, SD 14,187.1) relative to white-tailed deer (mean 17,828.4, min 4847, max 39,146, SD 7212.12). Age class and sex averages, in addition to winter data, are shown in Table S3. A Bray-Curtis PCoA clearly separated mountain goats and white-tailed deer (Fig. 2).

**Fig. 2.**
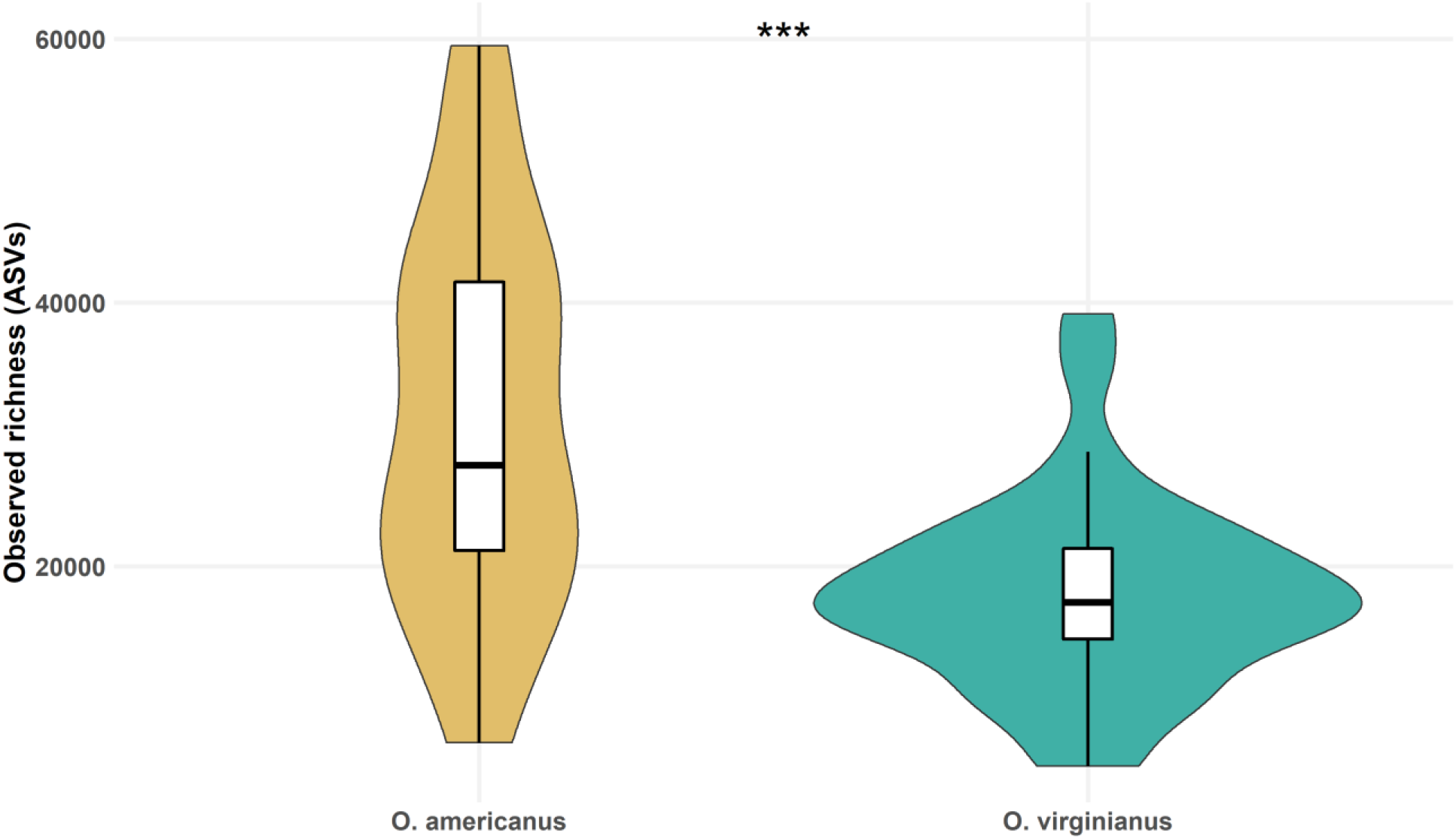
Observed richness (ASVs) for mountain goats (*Oreamnos americanus*, n=22) and white-tailed deer (*Odocoileus virginianus*, n=25). Mean observed richness was significantly different as computed by a two-sample t-test (t=4.11, df=29.64, p<0.001).

### Core range and proportional habitat use

Data filtering resulted in 84,932 GPS points for mountain goats (mean per individual 3,679, range 277 - 6,704, SD 761), and 63,900 GPS points for white-tailed deer (mean per individual 2,556, range 831 - 3,558, SD 906). One individual mountain goat was removed due to a small number of GPS points (n = 16). Mountain goats exhibited slightly larger summer core range size of 0.40 km^2^ (min 0.01 km^2^, max 0.72 km^2^, SD 0.17), compared to 0.36 km^2^ (min 0.13 km^2^, max 0.81 km^2^, SD 0.16) for white-tailed deer. The mean proportional habitat use values are reported in Table S4.

### Generalized linear models

We compared 50% isopleths from summer and winter; summer core ranges were reported here, as winter core ranges produced qualitatively similar results, albeit with weaker signals (Table S5). In both species, a greater *Firmicutes* to *Bacteroidetes* ratio was associated with larger core ranges with both models explaining an equal amount of variance (Nagelkerkes’s R^2^~0.28; Table 1; Fig. 3).

**Table 1.** Generalized linear models for core summer range size of mountain goat (*Oreamnos americanus*, n=22) and white-tailed deer (*Odocoileus virginianus*, n=25). The left column refers to the model where *Firmicutes* to *Bacteroides* ratios was a predictor, the middle column refers to the model where Pielou’s evenness was a predictor, and the right column refers to the model where observed richness (ASVs) was the predictor.

| Mountain goat |  |  |  |  |  |  |
| --- | --- | --- | --- | --- | --- | --- |
| Core range size |  |  |  |  |  |  |
| Predictors | Estimate | CI | Estimate | CI | Estimate | CI |
| (Intercept) | 0.31 | 0.10 - 0.51 | -2.3 | -4.1 - -0.42 | 0.19 | 0.026 - 0.35 |
| F:B Ratio | 0.010 | -0.019 - 0.038 | - | - | - | - |
| Pielou's evenness | - | - | 2.8 | 0.85 - 4.8 | - | - |
| Observed richness (ASVs) | - | - | - | - | $5.7 \times 10^{-6}$ | $1.4 \times 10^{-6} - 1.0 \times 10^{-5}$ |
| Age class (Subadult) | 0.20 | 0.018 - 0.39 | 0.17 | 0.020 - 0.32 | 0.16 | $3.3 \times 10^{-4} - 0.31$ |
| Sex (Male) | -0.030 | -0.17 - 0.10 | -<br>0.0047 | -0.12 - 0.11 | -0.012 | -0.13 - 0.11 |
| R<sup>2</sup> Nagelkerke | 0.28 |  | 0.47 |  | 0.45 |  |
| White-tailed deer |  |  |  |  |  |  |
| Core range size |  |  |  |  |  |  |
| Predictors | Estimate | CI | Estimate | CI | Estimate | CI |
| (Intercept) | 0.22 | 0.11 - 0.32 | 4.27 | -0.67 - 9.21 | 0.26 | 0.078 - 0.44 |
| F:B Ratio | 0.015 | 0.0050 - 0.025 | - | - | - | - |
| Pielou's evenness | - | - | -4.16 | -9.36 - 1.0 | - | - |
| Observed richness (ASVs) | - | - | - | - | $3.7 \times 10^{-6}$ | $-4.8 \times 10^{-6} - 1.2 \times 10^{-5}$ |
| Age class<br>(Subadult) | 0.033 | -0.084 - 0.15 | 0.061 | -0.068 - 0.19 | 0.069 | -0.064 - 0.20 |
| R<sup>2</sup> Nagelkerke | 0.28 |  | 0.21 |  | 0.075 |  |

**Fig. 3.**
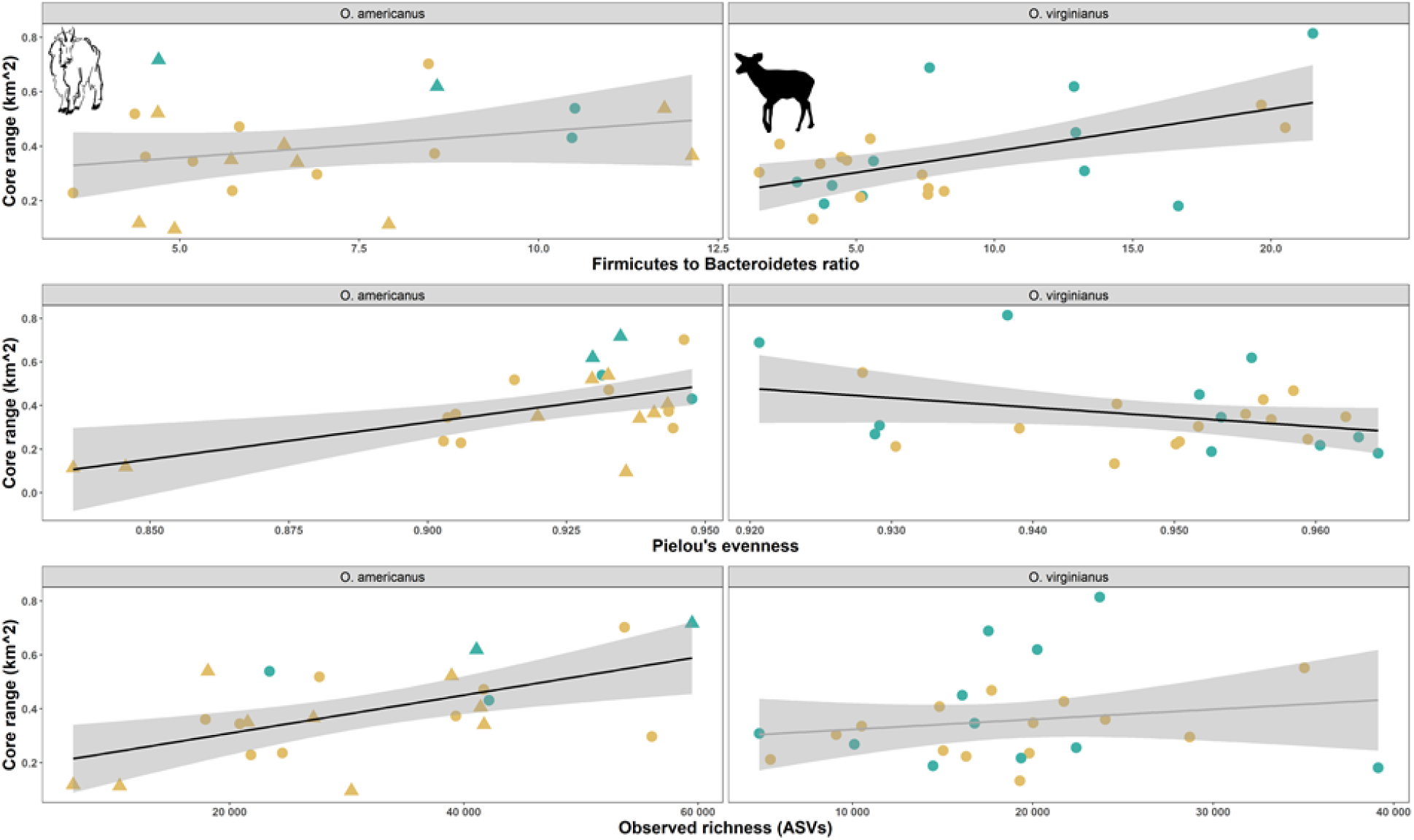
Microbiome metrics for mountain goats (*Oreamnos americanus*, n=22) northeast of Smithers, British Columbia, and white-tailed deer (*Odocoileus virginianus*, n=25) southwest of Ottawa, Ontario, relative to core summer range size. Females are represented by circles and only mountain goats have mixed-sex data. Adults (≥3 years of age) are represented in yellow, and subadults in blue. Significant Generalized Linear Models (GLMs) are denoted with a black line.

Pielou’s evenness was directly correlated with core range size in mountain goats; conversely, Pielou’s evenness was inversely correlated with core range size in white-tailed deer. Here the mountain goat model explained a relatively large portion of the variance (R^2^=0.47; Table 1; Fig. 3). Observed richness (ASVs) was associated with larger core ranges in mountain goats (Nagelkerkes’s R^2^~0.45, Table 1; Fig. 3) and white-tailed deer (Nagelkerkes’s R^2^~0.08, Table 1; Fig. 3). Age-class was a significant predictor in all three of the mountain goat core range GLMs, but neither of the white-tailed deer core range models while sex was a non-significant predictor in all mountain goat core range models (Table 1).

The use of escape terrain and treed areas was moderately correlated in mountain goats (t20=2.94, p<0.01, r=0.55), and exhibited significant relationships relative to Pielou’s evenness and number of ASVs; effect size confidence intervals did not overlap zero in models that measured the relationship between use of escape terrain and treed areas and Pielou’s evenness, and ASVs for the latter (Table 2). Specifically, a larger Pielou’s evenness value and number of ASVs was seen in individuals using less treed habitat and less escape terrain (Fig. 4). In HLI GLMs, confidence intervals overlapped zero and exhibited a slight decrease in R^2^ value relative to other mountain goat habitat use models (Table 2). All GLM parameter estimates in the white-tailed deer models had confidence intervals overlapping zero. Cross validation of linear models supported retaining age class and microbiome metrics as predictor variables of core range size. RMSE values in models with core range size as the response variable ranged from 0.12 to 0.18, and SI values ranged from 0.30 to 0.51, whereas in proportional habitat use models, RMSE values ranged from 0.11 to 0.33 and SI values from 0.033 to 1.14 (Table S6); this suggests moderate to high support for the models.

**Table 2.**
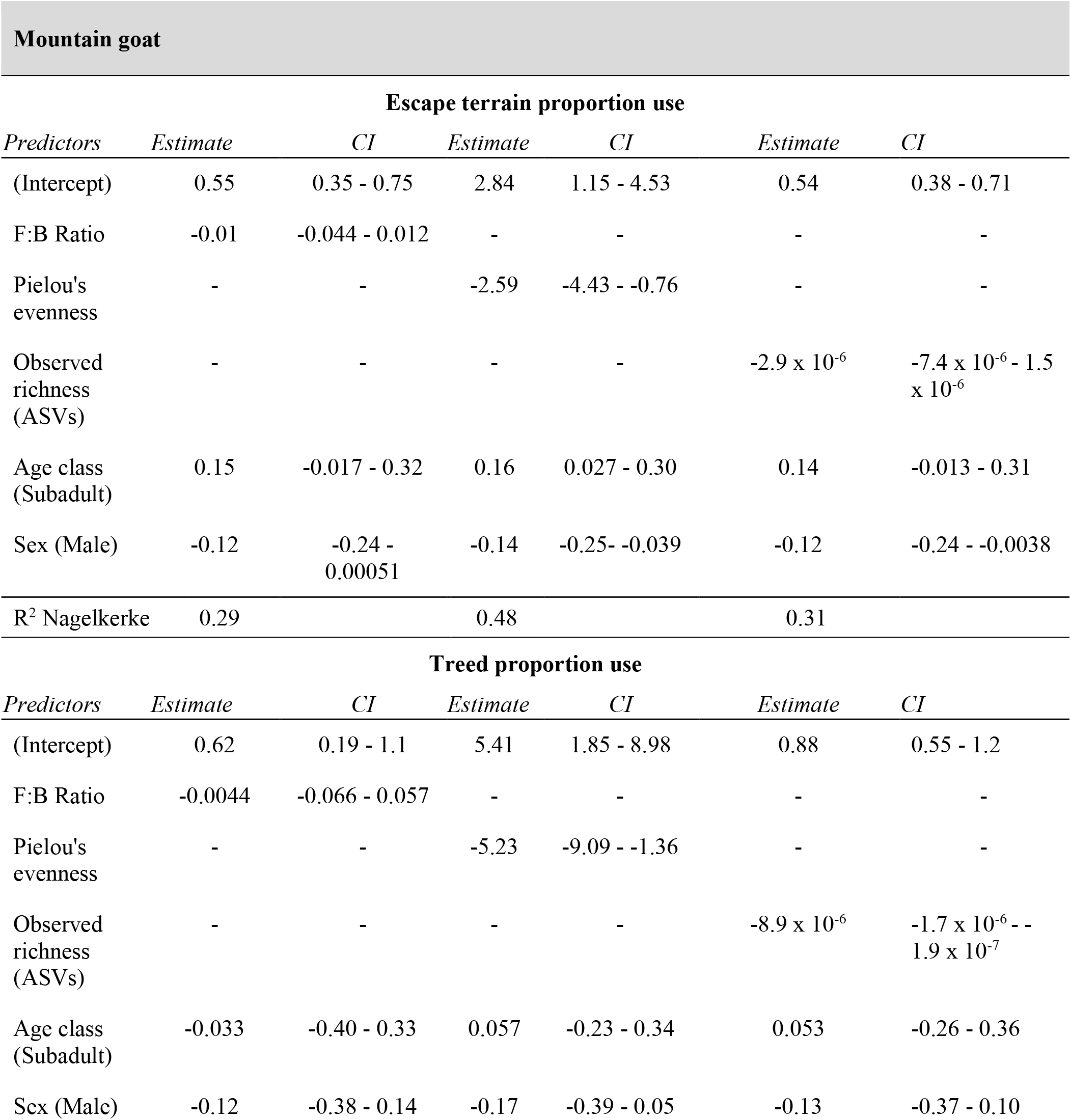
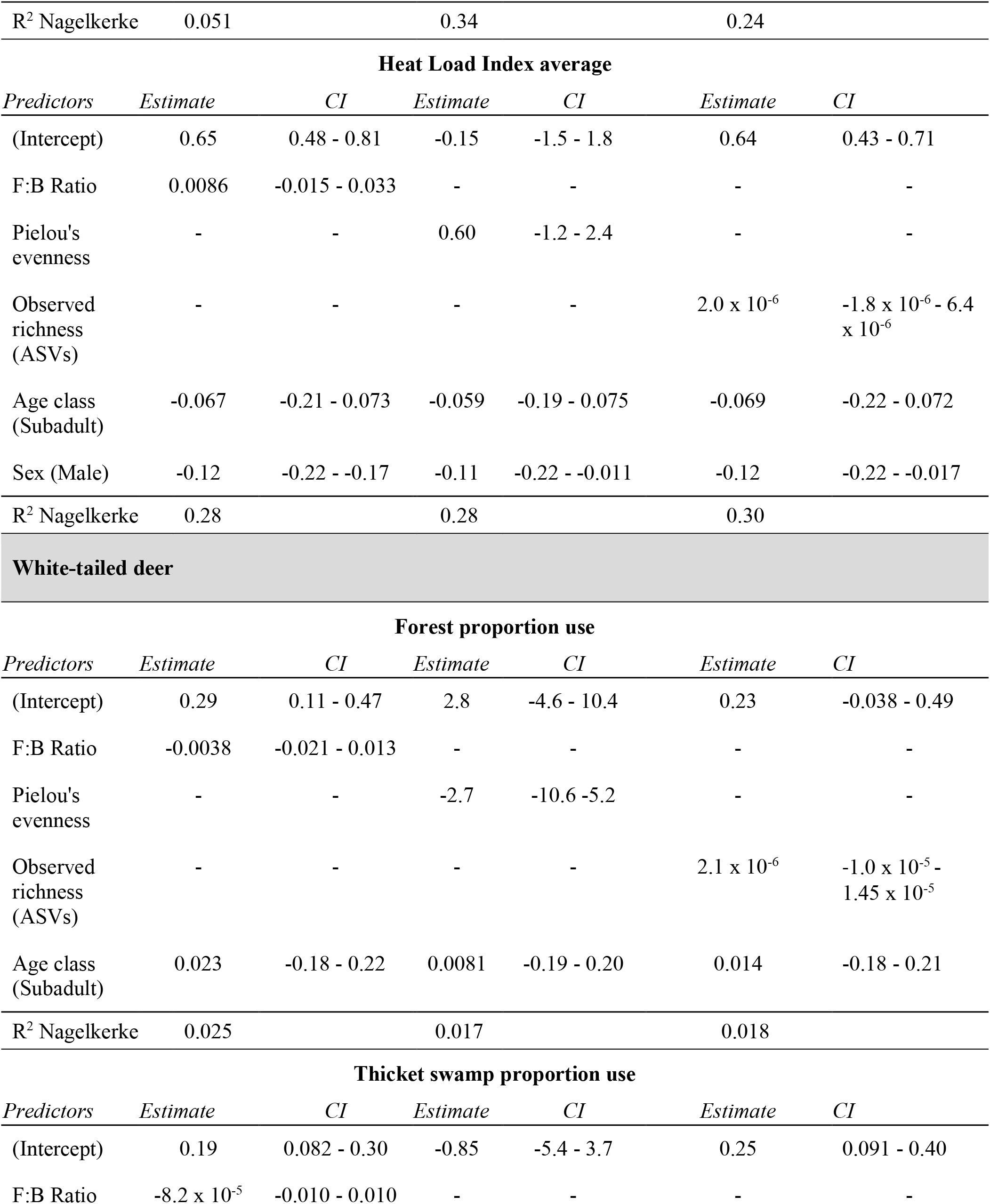
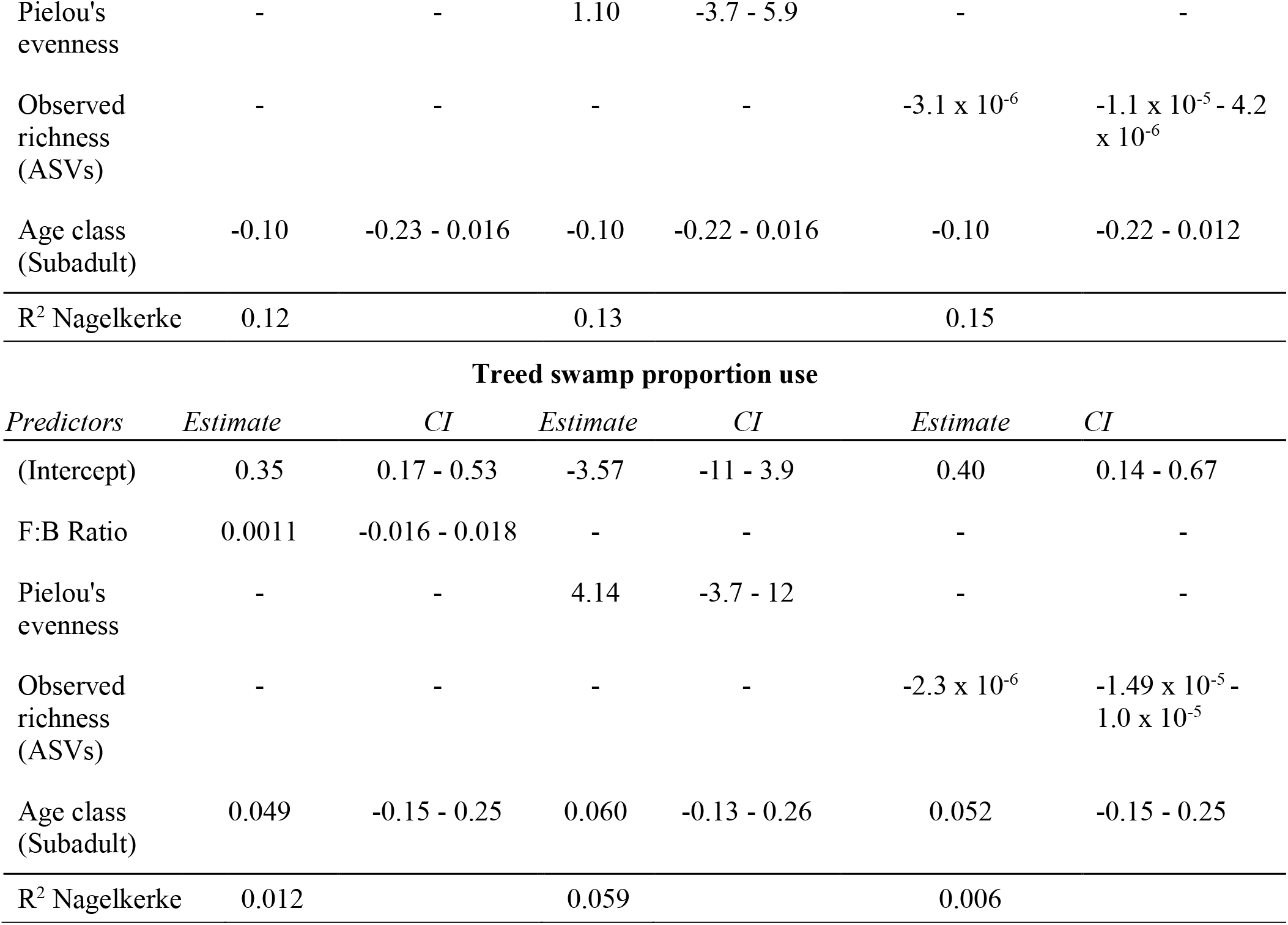
Generalized linear models for habitat proportion use for mountain goats (*Oreamnos americanus*, n=22) and white-tailed deer (*Odocoileus virginianus*, n=25). The left column refers to the model where *Firmicutes* to *Bacteroides* ratios was a predictor, the middle column refers to the model where Pielou’s evenness was a predictor, and the right column refers to the model where observed richness (ASVs) was the predictor.

| Mountain goat |  |  |  |  |  |  |
| --- | --- | --- | --- | --- | --- | --- |
| Escape terrain proportion use |  |  |  |  |  |  |
| Predictors | Estimate | CI | Estimate | CI | Estimate | CI |
| (Intercept) | 0.55 | 0.35 - 0.75 | 2.84 | 1.15 - 4.53 | 0.54 | 0.38 - 0.71 |
| F:B Ratio | -0.01 | -0.044 - 0.012 | - | - | - | - |
| Pielou's evenness | - | - | -2.59 | -4.43 - -0.76 | - | - |
| Observed richness (ASVs) | - | - | - | - | -2.9 x 10<sup>-6</sup> | -7.4 x 10<sup>-6</sup> - 1.5 x 10<sup>-6</sup> |
| Age class (Subadult) | 0.15 | -0.017 - 0.32 | 0.16 | 0.027 - 0.30 | 0.14 | -0.013 - 0.31 |
| Sex (Male) | -0.12 | -0.24 - 0.00051 | -0.14 | -0.25 - -0.039 | -0.12 | -0.24 - -0.0038 |
| R<sup>2</sup> Nagelkerke | 0.29 |  | 0.48 |  | 0.31 |  |
| Treed proportion use |  |  |  |  |  |  |
| Predictors | Estimate | CI | Estimate | CI | Estimate | CI |
| (Intercept) | 0.62 | 0.19 - 1.1 | 5.41 | 1.85 - 8.98 | 0.88 | 0.55 - 1.2 |
| F:B Ratio | -0.0044 | -0.066 - 0.057 | - | - | - | - |
| Pielou's evenness | - | - | -5.23 | -9.09 - -1.36 | - | - |
| Observed richness (ASVs) | - | - | - | - | -8.9 x 10<sup>-6</sup> | -1.7 x 10<sup>-6</sup> - -1.9 x 10<sup>-7</sup> |
| Age class (Subadult) | -0.033 | -0.40 - 0.33 | 0.057 | -0.23 - 0.34 | 0.053 | -0.26 - 0.36 |
| Sex (Male) | -0.12 | -0.38 - 0.14 | -0.17 | -0.39 - 0.05 | -0.13 | -0.37 - 0.10 |
| R<sup>2</sup> Nagelkerke | 0.051 |  | 0.34 |  | 0.24 |  |
| **Heat Load Index average** |  |  |  |  |  |  |
| *Predictors* | *Estimate* | *CI* | *Estimate* | *CI* | *Estimate* | *CI* |
| (Intercept) | 0.65 | 0.48 - 0.81 | -0.15 | -1.5 - 1.8 | 0.64 | 0.43 - 0.71 |
| F:B Ratio | 0.0086 | -0.015 - 0.033 | - | - | - | - |
| Pielou's evenness | - | - | 0.60 | -1.2 - 2.4 | - | - |
| Observed richness (ASVs) | - | - | - | - | 2.0 x 10<sup>-6</sup> | -1.8 x 10<sup>-6</sup> - 6.4 x 10<sup>-6</sup> |
| Age class (Subadult) | -0.067 | -0.21 - 0.073 | -0.059 | -0.19 - 0.075 | -0.069 | -0.22 - 0.072 |
| Sex (Male) | -0.12 | -0.22 - -0.17 | -0.11 | -0.22 - -0.011 | -0.12 | -0.22 - -0.017 |
| R<sup>2</sup> Nagelkerke | 0.28 |  | 0.28 |  | 0.30 |  |
| White-tailed deer |  |  |  |  |  |  |
| **Forest proportion use** |  |  |  |  |  |  |
| *Predictors* | *Estimate* | *CI* | *Estimate* | *CI* | *Estimate* | *CI* |
| (Intercept) | 0.29 | 0.11 - 0.47 | 2.8 | -4.6 - 10.4 | 0.23 | -0.038 - 0.49 |
| F:B Ratio | -0.0038 | -0.021 - 0.013 | - | - | - | - |
| Pielou's evenness | - | - | -2.7 | -10.6 - 5.2 | - | - |
| Observed richness (ASVs) | - | - | - | - | 2.1 x 10<sup>-6</sup> | -1.0 x 10<sup>-5</sup> - 1.45 x 10<sup>-5</sup> |
| Age class (Subadult) | 0.023 | -0.18 - 0.22 | 0.0081 | -0.19 - 0.20 | 0.014 | -0.18 - 0.21 |
| R<sup>2</sup> Nagelkerke | 0.025 |  | 0.017 |  | 0.018 |  |
| **Thicket swamp proportion use** |  |  |  |  |  |  |
| *Predictors* | *Estimate* | *CI* | *Estimate* | *CI* | *Estimate* | *CI* |
| (Intercept) | 0.19 | 0.082 - 0.30 | -0.85 | -5.4 - 3.7 | 0.25 | 0.091 - 0.40 |
| F:B Ratio | -8.2 x 10<sup>-5</sup> | -0.010 - 0.010 | - | - | - | - |
| Pielou's evenness | - | - | 1.10 | -3.7 - 5.9 | - | - |
| Observed richness (ASVs) | - | - | - | - | $-3.1 \times 10^{-6}$ | $-1.1 \times 10^{-5} - 4.2 \times 10^{-6}$ |
| Age class (Subadult) | -0.10 | -0.23 - 0.016 | -0.10 | -0.22 - 0.016 | -0.10 | -0.22 - 0.012 |
| R<sup>2</sup> Nagelkerke | 0.12 |  | 0.13 |  | 0.15 |  |

**Treed swamp proportion use**
| *Predictors* | *Estimate* | *CI* | *Estimate* | *CI* | *Estimate* | *CI* |
| --- | --- | --- | --- | --- | --- | --- |
| (Intercept) | 0.35 | 0.17 - 0.53 | -3.57 | -11 - 3.9 | 0.40 | 0.14 - 0.67 |
| F:B Ratio | 0.0011 | -0.016 - 0.018 | - | - | - | - |
| Pielou's evenness | - | - | 4.14 | -3.7 - 12 | - | - |
| Observed richness (ASVs) | - | - | - | - | $-2.3 \times 10^{-6}$ | $-1.49 \times 10^{-5} - 1.0 \times 10^{-5}$ |
| Age class (Subadult) | 0.049 | -0.15 - 0.25 | 0.060 | -0.13 - 0.26 | 0.052 | -0.15 - 0.25 |
| R<sup>2</sup> Nagelkerke | 0.012 |  | 0.059 |  | 0.006 |  |

**Fig. 4.**
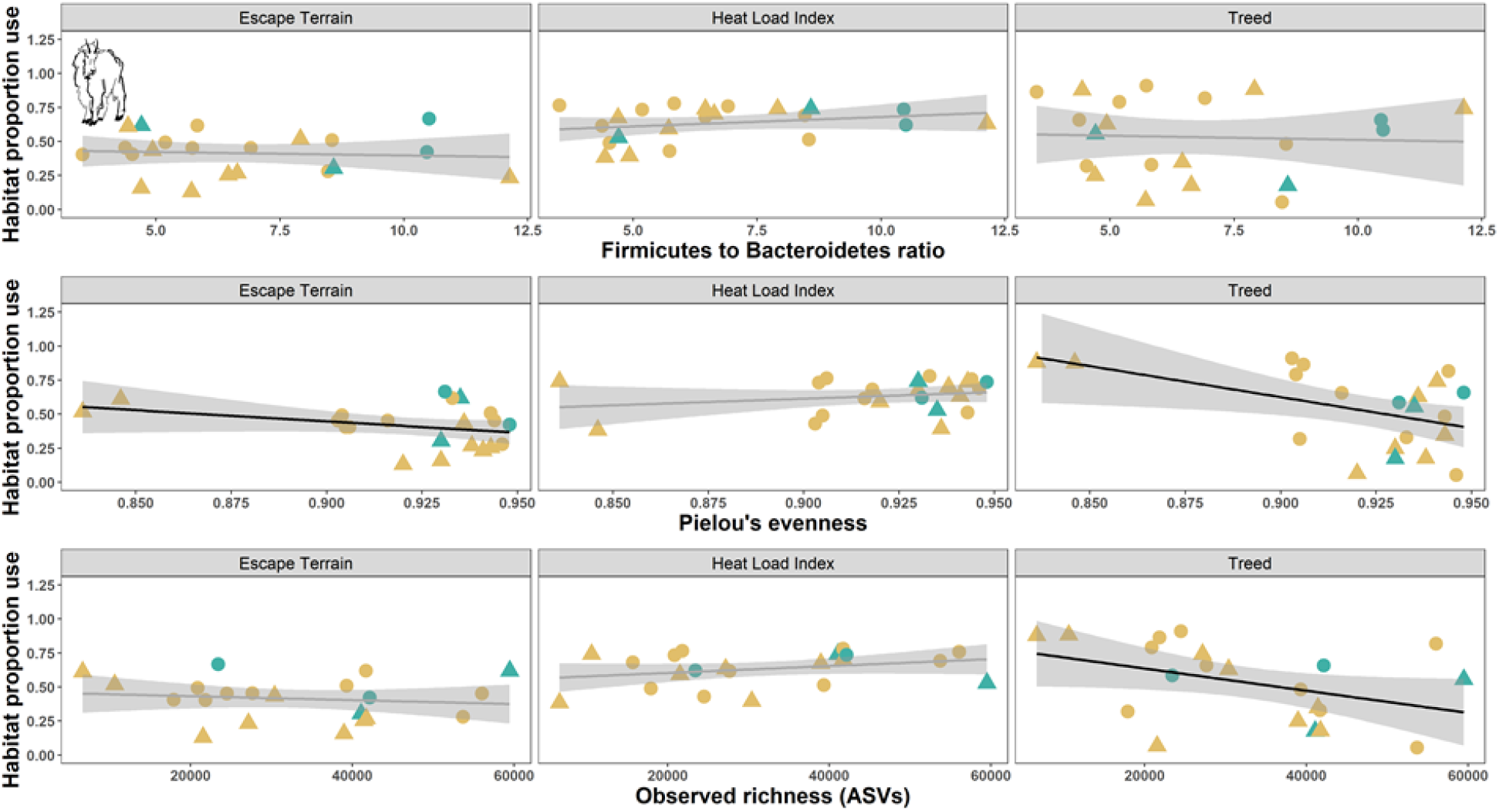
Microbiome metrics for mountain goats (*Oreamnos americanus*, n=21) northeast of Smithers, British Columbia, relative to habitat selection coefficients of both Heat Load Index (HLI) and treed habitat. Females are represented by circles and only mountain goats have mixed-sex data. Adults (≥3 years of age) are represented in yellow, and subadults in blue. Significant Generalized Linear Models (GLMs) are denoted with a black line.

## Discussion

The link between the gut microbiome and host space use has implications for foraging, activity levels, and ability to use energetically costly habitats. We show that certain aspects of the gut microbiome in both a generalist ungulate and specialist ungulate are linked to patterns of habitat use and home range size. Although the patterns are nuanced, there were some commonalities that collectively suggest the gut microbiota plays a role in determining space use patterns. An increase in *Firmicutes* to *Bacteroidetes* ratios in both species was correlated with an increase in core range sizes (Fig. 3). Increased *Firmicutes* to *Bacteroidetes* ratios are linked to increased Body Mass Index (BMI) and obesity in humans [19]. Both *Firmicutes* and *Bacteroidetes* are involved in energy resorption and are important for metabolism [62,63]. In a comparable study, home range size was not correlated with the *Firmicutes* to *Bacteroidetes* ratio in wild rodents [64], however, seasonal weight changes are more dynamic in small mammals [65,66], and we suspect this pattern might be more ubiquitous across temperate ungulates given their specialized gut microbiomes and need to put on fat stores. Ultimately, a higher *Firmicutes* to *Bacteroidetes* ratio may aid in the accumulation of fat stores to survive the winter.

In large ungulates, increasing levels of body fat is important to survive the winter when forage is limited, and may be impacted by the relative ratios of key bacterial species. In muskoxen (*Ovibos moschatus*), the abundance of *Firmicutes* stayed similar across seasons, while *Bacteroidetes* increased in the summer months, meaning the ratio of *Firmicutes* to *Bacteroidetes* is lower in the summer [67]. The ability to conserve bacteria necessary for adding fat, namely high levels of *Firmicutes*, while exhibiting changes in gut composition suggests ungulates can prepare for the winter even though the gut microbiome is shifting (see [39]). This concept is of importance to white-tailed deer, as diet turnover is pronounced in Ontario [68]. Shifting from herbaceous vegetation in the spring to woody browse in the winter may similarly result in increased diversity, while simultaneously conserving bacteria necessary for adding fat.

Specialists and generalists shift habitats seasonally, which can impact diet, and may be influenced by the gut microbiome; mountain goats move from alpine summer habitat to subalpine winter areas and white-tailed deer exhibit distinct winter and summer ranges [23,30,50]. There was considerably more variation in proportion of habitat used in the generalist white-tailed deer (Table S2), likely a function of their ability to use multiple habitat types and diet sources. Here, proportion habitat use-microbiome models had no relationship, as all proportion habitat use coefficients in white-tailed deer had confidence intervals overlapping zero (Table 2). Interestingly, the two habitat variables in mountain goats with clear signals were escape terrain and treed areas (Fig. 4), which are defining habitat characteristics of this species: the use of alpine terrain (i.e. no trees) and steep slopes [22,69]. Deviations from their specialized and preferred habitat types comes at a cost, as ungulates typically exhibit trade-offs with respect to forage quality and predation risk; avoiding predation can lead to decreased forage quality and abundance [70,71]. In mountain goats and bighorn sheep (*Ovis canadensis*), energy expenditures increased when travelling uphill or downhill - termed a vertical cost [72]. A lower *Firmicutes* to *Bacteroidetes* ratio could be linked to the vertical cost associated with spending more time in escape terrain, meaning less-fat or prime-conditioned individuals spend more time in escape terrain. Conversely, using more forage-available treed habitat may be correlated with an increase in *Firmicutes* to *Bacteroidetes* ratio as individuals have increased access to forage, with minimal vertical cost. Collectively, the habitat models had stronger signals in the specialist species compared to the generalist, and we suggest this is reflective of their more restricted habitat niche relative to generalists, ultimately supporting our predictions, where microbial deviations may be able to generate more prominent shifts in behaviour.

Deviations from the usage of important habitat types have larger consequences in specialists as they have a more restricted niche, and thus, the potential for the gut microbiome to modulate habitat use patterns is important in specialists to increase their competitive ability. While there is potential for modulation of proportion of specific habitat usage, the impact of the gut microbiome on the amount of space used may be more explicit. A lower Pielou’s evenness value indicates that a given individual has decreased diversity relative to another individual with the same number of gut microbial species, while a lower observed richness value represents a decrease in absolute number of gut microbiome species. The relationship between Pielou’s evenness and core range size differed in direction between species; greater gut evenness in mountain goats was correlated with a larger core range, while a negative relationship was noted in white-tailed deer (Fig. 3). Greater gut microbiome richness has been linked to increased gut stability in humans when faced with a shift in diet [73], as well as with changes in reproductive status in birds [74]. A positive relationship was noted between gut richness and core size in both species but was much stronger in mountain goats (Table 1; Fig. 3). Increased gut evenness might promote individuals moving through and foraging in larger areas, resulting in a positive relationship between evenness and core range size, while richness may act similarly. This is of added importance in a specialist, as the ability to utilize and forage in a more diverse array of habitat types might allow individuals to access and subsist in areas that other conspecifics cannot, thus increasing their competitive ability [75–77]. Higher relative gut microbiome diversity might allow specialists to forage more like generalists, while lower diversity may limit specialists to their typical niches. The reverse trend in white-tailed deer gut evenness might reflect the difference in feeding and habitat-use strategies; white-tailed deer are generalists, and thus by definition, can effectively use a variety of habitat and food sources. The negative relationship observed between gut evenness and range size in white-tailed deer might suggest that deer with higher gut microbiome evenness are able to use more diet sources overall and thus can meet their nutritional requirements within a smaller area relative to individuals with less diverse gut microbiomes. As generalists, deer are not as constrained by specific food and habitat types, and thus, their space use may be less heavily influenced by gut microbiome diversity. It is also possible that winter baiting had an impact on the gut microbiome in white-tailed deer, which could have led to weaker relationships.

We built our models assuming the microbiome influences habitat and home range patterns; but this need not be cause and effect, and the relationship is likely reciprocal [78]. For example, diet-microbiome covariance has been observed in multiple large herbivores, where seasonal diet turnover and seasonal microbiome turnover are positively correlated [79]. In this example, our model would assume the gut microbiome impacts diet choice (e.g. [11]) and in turn, the correlation between diet turnover and the microbiome contributes to habitat use of a given species. Assessing this relationship clearly needs experimental testing, and we view our study as a proof-of-concept that provides a testable hypothesis. Still, we demonstrated that quantifying the gut microbiome yields information related to space use; linking these two highly complex components of biology aids in our understanding of selection on the hologenome through the interplay between the individual and its microbial genomes and potential traits under selection (e.g. core range size and proportional habitat use). These findings also demonstrate that pellet sampling is useful in determining space and habitat use in managed populations, as it is conceivable with a large enough database and validation, one could predict the distribution and behaviour of animals on the landscape from non-invasively sampled pellets. Similar analyses of this kind should clarify the extent to which space use is linked to the gut microbiome in other species. Ultimately, we utilized a specialist and generalist ungulate to explore the link between the gut microbiome and movement to generate quantitative findings surrounding the impact of the gut microbiome on space use in wild populations.

## Acknowledgements

Special thanks to Jeff Bowman, Erin Koen, Kathleen Lo and Kiana Young for their comments on earlier drafts of this manuscript, Spencer Anderson for lab work, and Florent Déry for allowing us to use their digital drawings.

The authors declare that they have no conflict of interest.

## Data Accessibility

The sequence data generated for the mountain goat and white-tailed deer gut microbiota has been deposited in the Sequence Read Archive (BioProject ID: PRJNA638162; SRA submission: SUB7567698). R scripts used to perform analyses and generate figures are available at https://gitlab.com/WiDGeT_TrentU/graduate_theses/-/tree/master/Wolf_et_al.

## Author Contributions

J.F.W. and A.B.A.S. conceived the ideas; K.D.K., K.M.M., K.M., B.R.P., J.F.W., and A.B.A.S. collected materials and data; J.F.W. analysed the data; and J.F.W. wrote the paper with contributions from all co-authors.

