## Supplementary Information for "Gut microbiome composition predicts summer core range size in a generalist and specialist ungulate" (from another version of this preprint: version 2, 2020-08-05)

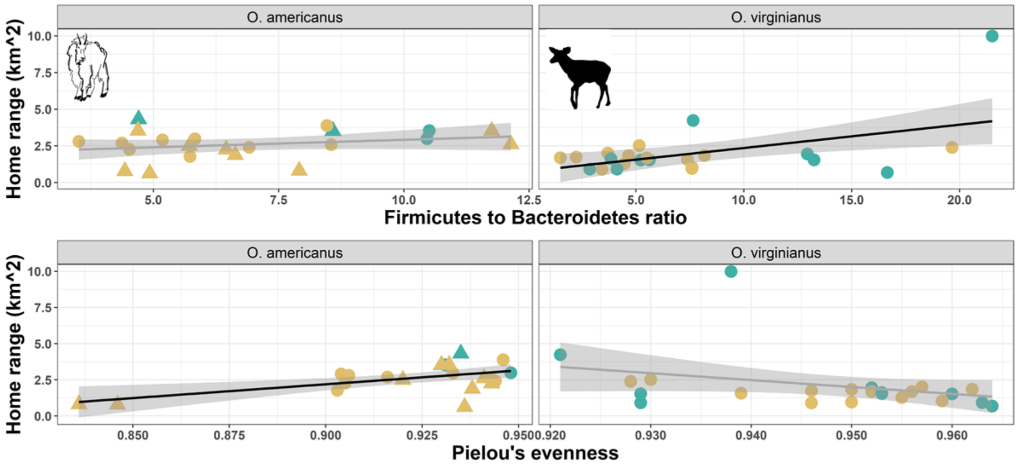


**Fig. S1** Microbiome metrics for mountain goats (*Oreamnos americanus*, n=22) northeast of Smithers, British Columbia, and white-tailed deer *(Odocoileus virginianus*, n=25) southwest of Ottawa, Ontario, relative to summer home range size. Females are represented by circles and only mountain goats have mixed-sex data. Adults (≥3 years of age) are represented in yellow, and subadults in blue. Significant Generalized Linear Models (GLMs) are denoted with a black line.

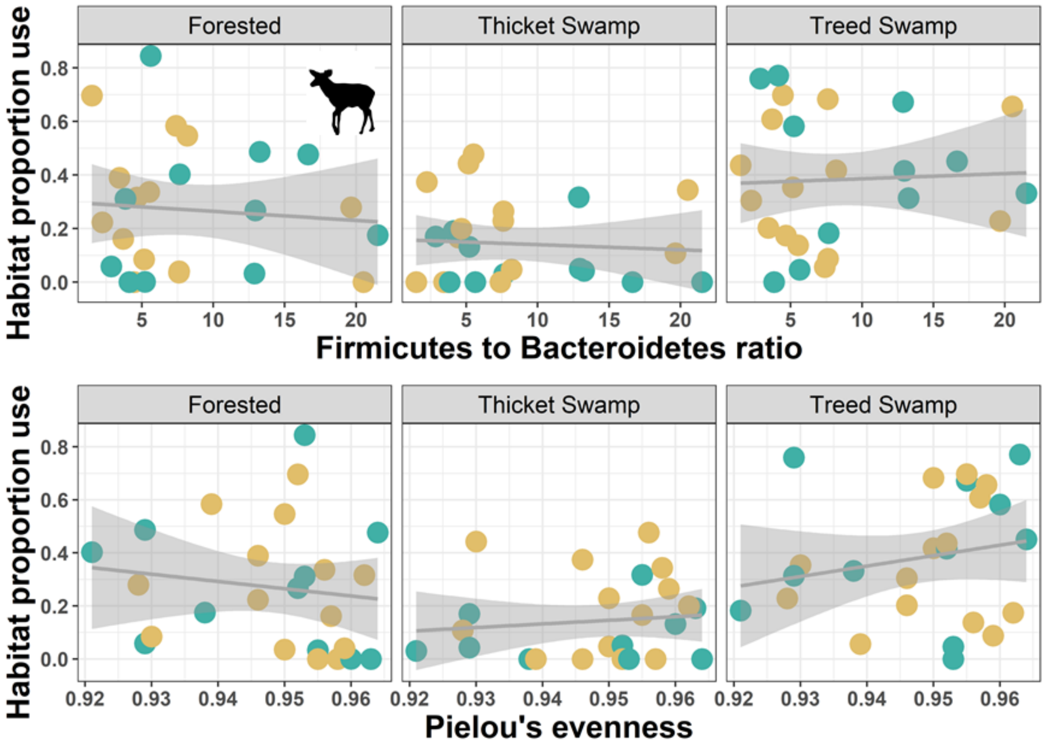


**Fig. S2** Microbiome metrics for white-tailed deer - *Odocoileus virginianus* southwest of Ottawa, Ontario, relative to forested, thicket swamp, and treed habitat proportion use (n=25). Adults (≥2 years of age) are represented in red, and subadults in blue. Significant Generalized Linear Models (GLMs) are denoted with a black line.

**Table S1** Metadata for habitat proportion use data among all mountain goat (*Oreamnos americanus*, n=22) samples used in the study

| Animal ID | 50% escape terrain proportion use | 95% escape terrain proportion use | 50% treed proportion use | 95% treed proportion use | 50% average Heat Load Index | 95% average Heat Load Index |
|---|---|---|---|---|---|---|
| MG1 | 0.249 | 0.155 | 0.158 | 0.098 | 0.675 | 0.655 |
| MG2 | 0.864 | 0.490 | 0.404 | 0.229 | 0.764 | 0.704 |
| MG3 | 0.791 | 0.473 | 0.492 | 0.294 | 0.732 | 0.684 |
| MG4 | 0.629 | 0.412 | 0.435 | 0.285 | 0.395 | 0.464 |
| MG5 | 0.910 | 0.551 | 0.453 | 0.274 | 0.430 | 0.467 |
| MG6 | 0.066 | 0.041 | 0.131 | 0.082 | 0.592 | 0.594 |
| MG8 | 0.053 | 0.033 | 0.279 | 0.173 | 0.681 | 0.674 |
| MG9 | 0.347 | 0.230 | 0.255 | 0.169 | 0.691 | 0.667 |
| MG10 | 0.318 | 0.186 | 0.406 | 0.237 | 0.738 | 0.692 |
| MG11 | 0.556 | 0.358 | 0.618 | 0.398 | 0.489 | 0.513 |
| MG12 | 0.175 | 0.109 | 0.303 | 0.189 | 0.529 | 0.535 |
| MG13 | 0.329 | 0.202 | 0.618 | 0.379 | 0.742 | 0.701 |
| MG14 | 0.482 | 0.291 | 0.508 | 0.306 | 0.780 | 0.715 |
| MG15 | 0.742 | 0.452 | 0.233 | 0.142 | 0.514 | 0.530 |
| MG16 | 0.658 | 0.412 | 0.453 | 0.284 | 0.632 | 0.620 |
| MG17 | 0.684 | 0.413 | 0.452 | 0.273 | 0.615 | 0.604 |
| MG18 | 0.585 | 0.360 | 0.666 | 0.409 | 0.620 | 0.581 |
| MG19 | N/A | N/A | N/A | N/A | N/A | N/A |
| MG20 | 0.177 | 0.118 | 0.268 | 0.179 | 0.702 | 0.683 |
| MG21 | 0.817 | 0.469 | 0.453 | 0.260 | 0.758 | 0.698 |
| MG22 | 0.658 | 0.391 | 0.423 | 0.251 | 0.736 | 0.688 |
| MG23 | 0.883 | 0.538 | 0.518 | 0.316 | 0.740 | 0.691 |
| MG24 | 0.249 | 0.155 | 0.158 | 0.098 | 0.384 | 0.409 |

**Table S2** Metadata for habitat proportion use data among all white-tailed deer (*Odocoileus virginianus*, n=25) samples used in the study

| Animal ID | 50% treed swamp proportion use | 95% treed swamp proportion use | 50% thicket swamp proportion use | 95% thicket swamp proportion use | 50% forest proportion use | 95% forest proportion use |
|---|---|---|---|---|---|---|
| D021a | 0.683 | 0.437 | 0.229 | 0.147 | 0.035 | 0.023 |
| D021b | 0.760 | 0.484 | 0.170 | 0.108 | 0.058 | 0.037 |
| D024a | 0.673 | 0.490 | 0.317 | 0.231 | 0.033 | 0.024 |
| D024b | 0.656 | 0.392 | 0.344 | 0.206 | 0.000 | 0.000 |
| D036 | 0.609 | 0.443 | 0.000 | 0.000 | 0.162 | 0.118 |
| D037a | 0.698 | 0.404 | 0.166 | 0.096 | 0.000 | 0.000 |
| D039 | 0.183 | 0.130 | 0.030 | 0.021 | 0.403 | 0.286 |
| D040 | 0.332 | 0.185 | 0.000 | 0.000 | 0.175 | 0.097 |
| D042 | 0.418 | 0.265 | 0.047 | 0.030 | 0.547 | 0.347 |
| D046 | 0.229 | 0.162 | 0.108 | 0.076 | 0.280 | 0.198 |
| D048 | 0.416 | 0.270 | 0.051 | 0.033 | 0.268 | 0.173 |
| D051 | 0.046 | 0.027 | 0.000 | 0.000 | 1.318 | 0.768 |
| D053 | 0.056 | 0.033 | 0.000 | 0.000 | 0.584 | 0.339 |
| D054 | 0.451 | 0.274 | 0.000 | 0.000 | 0.477 | 0.289 |
| D061 | 0.582 | 0.365 | 0.132 | 0.083 | 0.001 | 0.001 |
| D065 | 0.305 | 0.207 | 0.374 | 0.254 | 0.223 | 0.151 |
| D072 | 0.203 | 0.135 | 0.000 | 0.000 | 0.389 | 0.259 |
| D075 | 0.771 | 0.443 | 0.191 | 0.110 | 0.000 | 0.000 |
| D076 | 0.174 | 0.120 | 0.199 | 0.137 | 0.317 | 0.218 |
| D079 | 0.354 | 0.232 | 0.443 | 0.290 | 0.085 | 0.055 |
| D080 | 0.314 | 0.202 | 0.042 | 0.027 | 0.487 | 0.313 |
| D089 | 0.436 | 0.264 | 0.000 | 0.000 | 0.696 | 0.422 |
| D090 | 0.138 | 0.096 | 0.477 | 0.335 | 0.337 | 0.236 |
| D092 | 0.087 | 0.055 | 0.265 | 0.168 | 0.040 | 0.026 |
| D095 | 0.000 | 0.000 | 0.000 | 0.000 | 0.312 | 0.204 |

**Table S3** Metadata averages by sex and age class for gut microbiome and core range data among all mountain goats (*Oreamnos americanus*, n=23) and white-tailed deer (*Odocoileus virginianus*, n=25) samples used in the study

| Metric | Season | Age class | Sex |
|---|---|---|---|
| **Mountain goat** | | | |
| Pielou's evenness | | Adult - 0.92<br>Subadult - 0.94 | Male - 0.93<br>Female - 0.92 |
| *Firmicutes:Bacteroides* ratio | | Adult - 6.5<br>Subadult - 8.6 | Male - 7.1<br>Female - 6.7 |
| Core range size ($km^2$) | Winter | Adult - 0.057<br>Subadult -0.096 | Male - 0.051<br>Female - 0.076 |
| Core range size ($km^2$) | Summer | Adult -0.35<br>Subadult - 0.58 | Male - 0.38<br>Female - 0.41 |
| Escape terrain proportion use | Summer | Adult - 0.40<br>Subadult - 0.50 | Male - 0.35<br>Female- 0.47 |
| Treed selection proportion use | Summer | Adult - 0.55<br>Subadult - 0.49 | Male - 0.47<br>Female- 0.59 |
| Heat Load Index | Summer | Adult - 0.63<br>Subadult- 0.58 | Male - 0.57<br>Female - 0.66 |
| **White-tailed deer** | | | |
| Pielou's evenness | | Adult - 0.95<br>Subadult - 0.95 | |
| *Firmicutes:Bacteroides* ratio | | Adult - 6.2<br>Subadult - 9.4 | |
| Core range size ($km^2$) | Winter | Adult - 0.34<br>Subadult - 0.32 | |
| Core range size ($km^2$) | Summer | Adult - 0.31<br>Subadult - 0.37 | |
| Forest proportion use | Summer | Adult - 0.28<br>Subadult - 0.30 | |
| Treed swamp proportion use | Summer | Adult - 0.34<br>Subadult - 0.39 | |
| Thicket swamp proportion use | Summer | Adult - 0.18<br>Subadult - 0.06 | |

**Table S4** Average and standard deviations of habitat proportion use data among all mountain goat (*Oreamnos americanus*, n=22) and white-tailed deer (*Odocoileus virginianus*, n=25) samples used in the study

| Average | Standard deviation | Habitat type |
|---|---|---|
| **Mountain goat** | | |
| 0.54 | 0.28 | Treed |
| 0.41 | 0.15 | Escape terrain |
| 0.63 | 0.12 | Heat Load Index |
| **White-tailed deer** | | |
| 0.33 | 0.30 | Forested |
| 0.13 | 0.16 | Thicket swamp |
| 0.34 | 0.22 | Treed swamp |

**Table S5** Root Mean Square Error (RMSE) and Scatter Index (SI) values from five-fold linear model cross validation for all Generalized Linear Models (GLMs) reported in this study

| Model | RMSE | SI |
|---|---|---|
| **Mountain goat** | | |
| Core range ~ F:B + age class | 0.17 | 0.43 |
| Core range ~ Pielou + age class | 0.12 | 0.30 |
| Escape proportion use ~ F:B + age class | 0.15 | 0.37 |
| Escape proportion use ~ Pielou + age class | 0.13 | 0.32 |
| Treed proportion use ~ F:B + age class | 0.31 | 0.58 |
| Treed proportion use ~ Pielou + age class | 0.25 | 0.47 |
| Heat Load Index proportion use ~ F:B + age class | 0.11 | 0.18 |
| Heat Load Index proportion use ~ Pielou + age class | 0.11 | 0.18 |
| **White-tailed deer** | | |
| Core range ~ F:B + age class | 0.15 | 0.42 |
| Core range ~ Pielou + age class | 0.18 | 0.51 |
| Forest proportion use ~ F:B + age class | 0.33 | 1.14 |
| Forest proportion use ~ Pielou + age class | 0.31 | 1.09 |
| Thicket swamp proportion use ~ F:B + age class | 0.15 | 1.06 |
| Thicket swamp proportion use ~ Pielou + age class | 0.16 | 1.14 |
| Treed swamp proportion use ~ F:B + age class | 0.26 | 0.67 |
| Treed swamp proportion use ~ Pielou + age class | 0.24 | 0.63 |

**Table S6** Generalized linear models for **winter core range** size of mountain goats - *Oreamnos americanus* and white-tailed deer - *Odocoileus virginianus* ($n_{MG}$=22, $n_{WTD}$=19). The further left column refers to the model where *Firmicutes* to *Bacteroides* ratios was a predictor, while the right column refers to the model where Pielou’s evenness was a predictor

**Mountain goat**

| | **Core range size** | | **Core range size** | |
|---|---|---|---|---|
| *Predictors* | *Estimates* | *CI* | *Estimates* | *CI* |
| (Intercept) | 0.04 | -0.01 – 0.10 | -0.31 | -0.88 – 0.25 |
| F:B Ratio | 0.00 | -0.01 – 0.01 | - | - |
| Pielou's evenness | - | -- | 0.40 | -0.21 – 1.02 |
| Age class (Subadult) | 0.04 | -0.01 – 0.08 | 0.03 | -0.01 – 0.08 |
| $R^2$ Nagelkerke | 0.14 | | 0.19 | |

**White-tailed deer**

| | **Core range size** | | **Core range size** | |
|---|---|---|---|---|
| *Predictors* | *Estimates* | *CI* | *Estimates* | *CI* |
| (Intercept) | 0.42 | 0.25 – 0.58 | -1.98 | -9.53 – 5.58 |
| F:B Ratio | -0.01 | -0.03 – 0.01 | - | - |
| Pielou's evenness | - | - | 2.45 | -5.50 – 10.39 |
| Age class (Subadult) | -0.00 | -0.20 – 0.19 | -0.04 | -0.23 – 0.14 |
| $R^2$ Nagelkerke | 0.088 | | 0.036 | |
